# Identification of alcohol use disorder-associated and ethanol-responsive proteins and candidate biomarkers using human cortical organoids

**DOI:** 10.64898/2026.09.27.754852

**Authors:** Xindi Li, Andrew J. Boreland, Arpana Agrawal, Chella Kamarajan, Paul A. Slesinger, Jay A. Tischfield, Yue Wang, Howard J. Edenberg, Tatiana Foroud, Yunlong Liu, Jubao Duan, Ronald P. Hart, Zhiping P. Pang, Dongbing Lai

## Abstract

Genome-wide association studies of alcohol use disorder (AUD) have identified >100 loci, yet the functional impact and the downstream molecular mechanisms are not well understood. As functional effectors of genes, proteins provide direct insight into disease mechanisms. However, most proteomic alterations in postmortem human brains of AUD likely reflect a combination of inherited susceptibility (AUD-associated), the consequences of alcohol exposure (ethanol-responsive), other environmental and lifestyle factors, as well as sample preparation. Here, we profiled the proteome of human microglia-containing cortical organoids derived from human induced pluripotent stem cells from individuals with AUD (n=11) and without (n=5), paired with or without intermittent ethanol exposure. To maximize statistical power, donors were selected based on high or low genetic liability for AUD as measured by polygenic scores. Among 8,952 proteins that passed quality control, 1,038 were AUD-associated and 718 were ethanol-responsive. AUD-associated proteins were enriched for neuronal signaling, immune, mitochondrial, and extracellular matrix pathways, whereas ethanol-responsive proteins were enriched for RNA processing, ribosome biogenesis, protein translation, vesicle trafficking, and membrane transport pathways. Twenty ethanol-responsive proteins were replicated in a postmortem human prefrontal cortex proteomic dataset. Using UK Biobank plasma proteomics data, we identified 26 candidate AUD-associated biomarkers and 9 candidate ethanol-responsive biomarkers. We also found that major proteomic variation was significantly correlated with AUD polygenic scores. Together, these findings reveal distinct proteomic signatures of inherited susceptibility to AUD and ethanol exposure, providing new insights into mechanisms underlying AUD and identifying candidate circulating biomarkers for further investigation.

## Introduction

Alcohol use disorder (AUD), characterized by excessive and uncontrolled alcohol consumption despite adverse social, mental, and health consequences, represents a significant public health challenge^1^. In the U.S., the 2025 National Survey on Drug Use and Health indicates that 25.7 million people aged ≥12 years had AUD in the past year^2^. Besides the direct adverse consequences, AUD significantly elevates the likelihood for >200 mental and physical health problems and diseases^3^. Given its profound individual, familial, and societal impact, elucidating the genetic and molecular mechanisms underlying AUD is essential for developing novel and more effective strategies for its prevention and treatment.

AUD is a highly polygenic psychiatric disorder^4–6^, yet the molecular mechanisms linking genetic susceptibility to disease pathophysiology remain poorly understood. As the functional products of genes, proteins provide direct insight into potential etiological processes underlying AUD by revealing dysregulated biological pathways, while also enabling the identification of circulating biomarkers for diagnosis/prognosis and potential targets for therapeutic development. Given their psychiatric basis, studying brain-based proteomic process is of particular utility to AUD research. However, proteomic studies of postmortem human brain tissue are complicated by substantial biological heterogeneity, because brain proteomes are influenced by age, lifestyle, comorbid conditions, and the cumulative effects of chronic alcohol exposure^7, 8^, as well as technical factors such as postmortem interval (PMI), tissue pH, and sample processing^9^.

Although previous studies have matched cases and controls on selected demographic and clinical characteristics^8, 10–18^, it is difficult to adequately control for these and other potential confounders. Several studies have integrated large-scale GWAS with human brain proteomic data using Mendelian randomization to prioritize causal proteins; however, these analyses were restricted to proteins linked to genome-wide significant variants and relied on reference brain proteomes that were not derived from individuals with AUD^19–21^. Other studies have identified circulating protein biomarkers associated with AUD and related phenotypes^22–26^, but whether these plasma proteins reflect molecular alterations in the brain remains unclear.

Proteomic alterations observed in individuals with AUD may arise from two biologically distinct processes: inherited susceptibility to AUD (AUD-associated) and the downstream consequences of ethanol exposure (ethanol-responsive). AUD-associated proteins may reflect mechanisms that predispose individuals to the disorder and therefore represent potential prevention and therapeutic targets, whereas ethanol-responsive proteins may provide insight into the molecular adaptation consequent to prolonged alcohol intake and alcohol-related comorbidities, and may help with the management of symptoms and increasing the likelihood of remission. These processes are difficult to disentangle in postmortem human brain tissue because samples from individuals with AUD are obtained only after years of alcohol exposure.

Brain organoids derived from human induced pluripotent stem cells (hiPSCs) retain the donor’s genetic background therefore can be used to identify both AUD-associated and ethanol-responsive proteins^27^. However, there is no human brain organoid study that stratified donors by AUD status and genetic liability to AUD to identify AUD-associated proteins. A previous study exposed dorsal forebrain organoids to 100 mM ethanol for 7 days and identified ethanol-responsive changes in nervous system development, cellular stress responses, nonsense-mediated mRNA decay, and metabolites^28^. However, that study used organoids lacking microglia, which have been shown to contribute to AUD-related brain dysfunction in our previous studies^29, 30^.

To address these gaps, we utilized microglia-containing cortical organoids (mCOs) derived from donors with/without AUD alongside information on their genic liability as measured by polygenic scores (PGS). mCO from each donor were subjected to intermittent ethanol exposure in a paired design, enabling us to distinguish AUD-associated proteins from ethanol-responsive proteins while reducing inter-individual variability. Pathway and network analyses were performed to characterize the biological processes underlying these two types of proteins. We replicated ethanol-responsive proteins using proteomics data from human post-mortem prefrontal cortex (PFC) tissues. We also integrated our findings with UK Biobank plasma proteomic data to identify candidate circulating biomarkers. We further assessed the relationship between major proteomic variation and genetic liability for AUD, as measured by polygenic scores.

## Methods

### Human microglia-containing cortical organoids (mCOs)

Neural progenitor cells (NPCs) and primitive macrophage precursors (PMPs) were independently differentiated from the same Human induced pluripotent stem cell (hiPSC) donors and then combined to form donor-matched mCOs. hiPSC lines were derived from participants in the Collaborative Study on the Genetics of Alcoholism (COGA^31–33^: 9 males and 7 females, all European ancestry), which are available from the COGA Sharing Repository (https://cogastudy.org/resources-for-researchers/). Eleven donors met criteria for a lifetime history of AUD, and their PGS for AUD were among top 10% in COGA^34^, and 5 donors did not meet criteria for AUD, and their PGS for AUD were among bottom 10% in COGA^34^. Donors were selected using the PGS derived by Barr and colleagues^34^ (hereafter referred to as the original PGS). Because PGS performance can vary depending on the discovery GWAS used for their derivation, we also recalculated PGS using the largest GWAS of problematic alcohol use (PAU)^51^ (hereafter referred to as the latest PGS) to assess the consistency of the findings across different measures of PGS.

Human primitive neural progenitor cells (pNPCs) were generated from hiPSCs by dual-SMAD inhibition using SB431542 (5 µM; Stemgent) and Noggin (50 ng/mL; PeproTech) in neural induction medium consisting of DMEM/F12 (HyClone) supplemented with 1× N2 (Thermo Fisher Scientific) for 1 week. Neural rosettes were then isolated and expanded in pNPC medium consisting of a 1:1 mixture of Neurobasal medium and DMEM/F12 supplemented with 1× N2, 1× B27 without retinoic acid, FGF2 (20 ng/mL; PeproTech), human leukemia inhibitory factor (hLIF; 10 ng/mL; Millipore), CHIR99021 (3 µM; Biogems), and SB431542 (2 µM), as previously described^35, 36^. Human PMPs were prepared from hiPSCs using a previously described protocol^37^.

For mCO generation, we aggregated pNPCs at 10,000 cells per well in ultra-low-attachment plates. After 2 weeks, mCOs were transferred to an orbital shaker and maintained in neuronal differentiation medium consisting of a 1:1 mixture of Neurobasal medium and DMEM/F12 supplemented with 1× N2, 1× B27, NT-3 (10 ng/ml; PeproTech), BDNF (10 ng/ml; PeproTech), GDNF (10 ng/ml; PeproTech), dibutyryl-cAMP (1 mM; Sigma), and ascorbic acid (200 nM; Sigma). The medium was replaced every other day. On day 30, 2 × 10^4 PMPs were added to each mCO, and the resulting mCOs were maintained in neuronal differentiation medium supplemented with interleukin-34 (IL-34) (100 ng/ml; PeproTech, catalog #200-34) and granulocyte-macrophage colony-stimulating factor (GM-CSF) (10 ng/ml; PeproTech, catalog #300-03).

### Intermittent ethanol exposure

On day 90, a 10-day protocol of intermittent ethanol exposure was performed as previously described^29, 30, 38^. On the first day of treatment, all medium was removed from each well and replaced with neuronal differentiation medium supplemented with ethanol (200 proof, non-denatured; Decon Laboratories) to achieve peak concentrations of 0 or 75 mM. Accounting for the rate of evaporation, this exposure paradigm produced a mean 24-hour ethanol concentration of 28.79 mM, corresponding to a blood alcohol concentration of approximately 0.13%^38^.

On the 2^nd^, 3^rd^, 5^th^, 6^th^, 8^th^, and 9^th^ days of treatment, half of the medium was removed from each well, ethanol was added to the removed medium at twice the 75 mM concentration, and the mixture was returned to the cultures. On 4^th^ and 7^th^ days, a half-medium change was performed by replacing half of the existing medium with fresh medium containing ethanol at twice the 75 mM concentration. On day 100, cultures were collected for LC-MS.

### Proteomics data generation

Sample preparation and LC-MS were conducted by MetwareBio (Woburn, MA). Proteomic data were acquired using data-independent acquisition (DIA) and the resulting data were analyzed using DIA-NN (v1.8.1) with the library-free method^39^. The UP000005640_human_20230504.fasta database (A total of 82492 sequences) was used to create a spectra library with deep learning algorithms of neural networks. The option of MBR (Match Between Runs) was employed to create a spectral library from DIA data and then reanalyzed using this library. False discovery rate (FDR) of search results was adjusted to <1% at both protein and precursor ion levels, the remaining identifications were used for further quantification analysis.

Because missing values in LC-MS proteomics may reflect either protein not expressed or technical factors, we restricted downstream analyses to proteins with missing in ≤20% of samples (i.e., missing in ≤6 samples). Protein abundances were normalized within samples to generate relative protein abundances that were comparable across samples. The normalized protein abundance data was used for principal component analysis (PCA) to identify major sources of biological variation, including AUD status, ethanol exposure, and PGS, as well as potential technical variation, and for subsequent identification of AUD-associated and ethanol-responsive proteins. All statistical analyses were performed using R (v4.4.1).

### Identification of AUD-associated and ethanol-responsive proteins

To identify AUD-associated proteins, generalized linear mixed-effects (GLMM) models with a binomial distribution and logit link were fitted with AUD status as the dependent variable, normalized protein abundances as the independent variable, ethanol treatment as a fixed-effect covariate, and donor as a random effect to account for the paired untreated and ethanol-exposed samples. To identify ethanol-responsive proteins, paired t-tests were performed comparing ethanol-treated and untreated mCOs from the same donor to isolate the effect of ethanol exposure. Because this study was designed as an exploratory discovery analysis to prioritize candidate proteins for downstream pathway and network analyses, and because AUD is a highly polygenic disorder due to perturbations of many gene products, statistical significance was defined as FDR <0.20 to maximize sensitivity while maintaining control of multiple testing.

### Gene Set Enrichment Analysis (GSEA) using Reactome pathways

GSEA was performed using the fgsea R package^40, 41^ on pre-ranked lists of proteins. GSEA evaluates proteome-wide pathway enrichment, therefore, all quantified proteins were retained rather than restricting the analysis to differentially expressed proteins using a predefined significance threshold. Proteins were ranked according to their Z-scores (GLMM) or t-statistics (paired t-test), with positive and negative values indicating increased and decreased AUD likelihood or protein abundances. To reduce the influence of extreme values, Z-scores were capped at 20 before ranking. Reactome pathways were obtained using the msigdbr R package^40, 42^. GSEA was performed with a protein set size between 10 and 500, and eps = 0.0 to improve the accuracy of small P-value estimation. Statistical significance was assessed using the normalized enrichment score (NES), and P-values were adjusted for multiple testing using the Benjamini–Hochberg method^43^. Pathways with an FDR <0.05 were considered significantly enriched as suggested^40, 41^.

### Disease enrichment analyses

To assess whether identified proteins were associated with other diseases, including potential comorbidities, we performed disease enrichment using the DOSE R package^44^ with the DisGeNET database^45^. Unlike GSEA, which uses a ranked list of all proteins, disease enrichment analysis is an overrepresentation analysis that requires a predefined list of significant proteins. Therefore, proteins meeting the significance criteria (FDR <0.20) were included. Three disease enrichment analyses were performed separately: (1) proteins associated with increased likelihood of AUD; (2) ethanol-responsive proteins with increased abundances following ethanol treatment; and (3) ethanol-responsive proteins with decreased abundances following ethanol treatment. Separate analyses were conducted because combining proteins with opposite directions of association or abundance could obscure disease-specific enrichment patterns and reduce biological interpretability.

### Replication of ethanol-responsive proteins using proteomics data from human post-mortem prefrontal cortex (PFC) tissues

PFC proteomic data, performed using multiplex LC–MS, were obtained from Teng et al.^8^ and is publicly available through the ProteomeXchange Consortium^46^. To minimize potential population stratification, we included only individuals of genetically determined European ancestry in the analysis, which comprised 8 individuals with AUD and 13 matched controls (all males). A total of 4,706 proteins were quantified in both mCOs and postmortem PFC and were included in the replication analysis.

Because postmortem brain tissues include the cumulative effects of chronic alcohol exposure, replication analyses were restricted to ethanol-responsive proteins. We reperformed analyses using linear regression models with protein abundance as the dependent variable and daily alcohol intake at death (DAI) as the independent variable. PCA of the PFC proteomic data revealed a pronounced batch effect along the first principal component (PC1) and three distinct clusters along the second principal component (PC2); therefore, batch and PC2 were included as covariates. Proteins with P-value <0.05 and having the concordant effect were considered as replicated. To evaluate whether the observed replication rate exceeded that expected by chance, we performed a binomial test using Bin(N, 0.025), where N was the number of significant proteins identified in mCOs and was quantified in postmortem PFC, and 0.025 is the probability of having a P-value <0.05 and a concordant effect.

### Identification of candidate AUD-associated and ethanol-responsive biomarkers

Plasma proteomic data were from the UK Biobank Pharma Proteomics Project (UKB-PPP)^47^, which were generated using Olink Explore™ Proximity Extension Assay^47^, and a total of 1,465 proteins were quantified in both UKB-PPP and mCOs. Proteins associated with incident AUD were considered as candidate AUD-associated biomarkers as AUD developed after plasma collection; proteins associated with prevalent AUD were considered as candidate ethanol-responsive biomarkers as plasma were collected after prolonged alcohol exposure. Proteins associated with incident and prevalent AUD were identified by Deng and colleagues^22^, which were downloaded from https://proteome-phenome-atlas.com/. The same binomial test as in replication analysis was used to evaluate whether the rates of identified biomarkers exceeded that expected by chance.

### Differential ethanol responses in AUD and non-AUD organoids using weighted gene co-expression network analysis (WGCNA)

Protein abundance data were log₂-transformed. A signed network was constructed using biweight midcorrelation^48^. We selected the soft-thresholding power based on the scale-free topology, using a default of 6 when no suitable value was identified. Modules were detected using a signed topological overlap matrix, a minimum module size of 20, and a merge threshold of 0.25. For each module, the module eigengene^49^, defined as the first principal component summarizing the protein abundance pattern within that module, was used as the dependent variable in a linear mixed-effects model. AUD status, ethanol exposure, and their interaction were included as fixed effects, with donor included as a random effect. Ethanol effects within the AUD and non-AUD groups, as well as the AUD and ethanol interaction, were estimated from the fitted models.

## Results

We derived hiPSCs from 16 individuals with or without a lifetime history of AUD who were drawn from the top 10% and bottom 10% of the distribution of PGS within COGA^30^. hiPSCs were differentiated into microglia-containing cortical brain organoids; then underwent a 10-day intermittent exposure to a mean ethanol concentration of 28.79 mM over 24 hours or were cultured without ethanol followed by proteomics analyses (**Figure 1A**).

**Figure 1.**
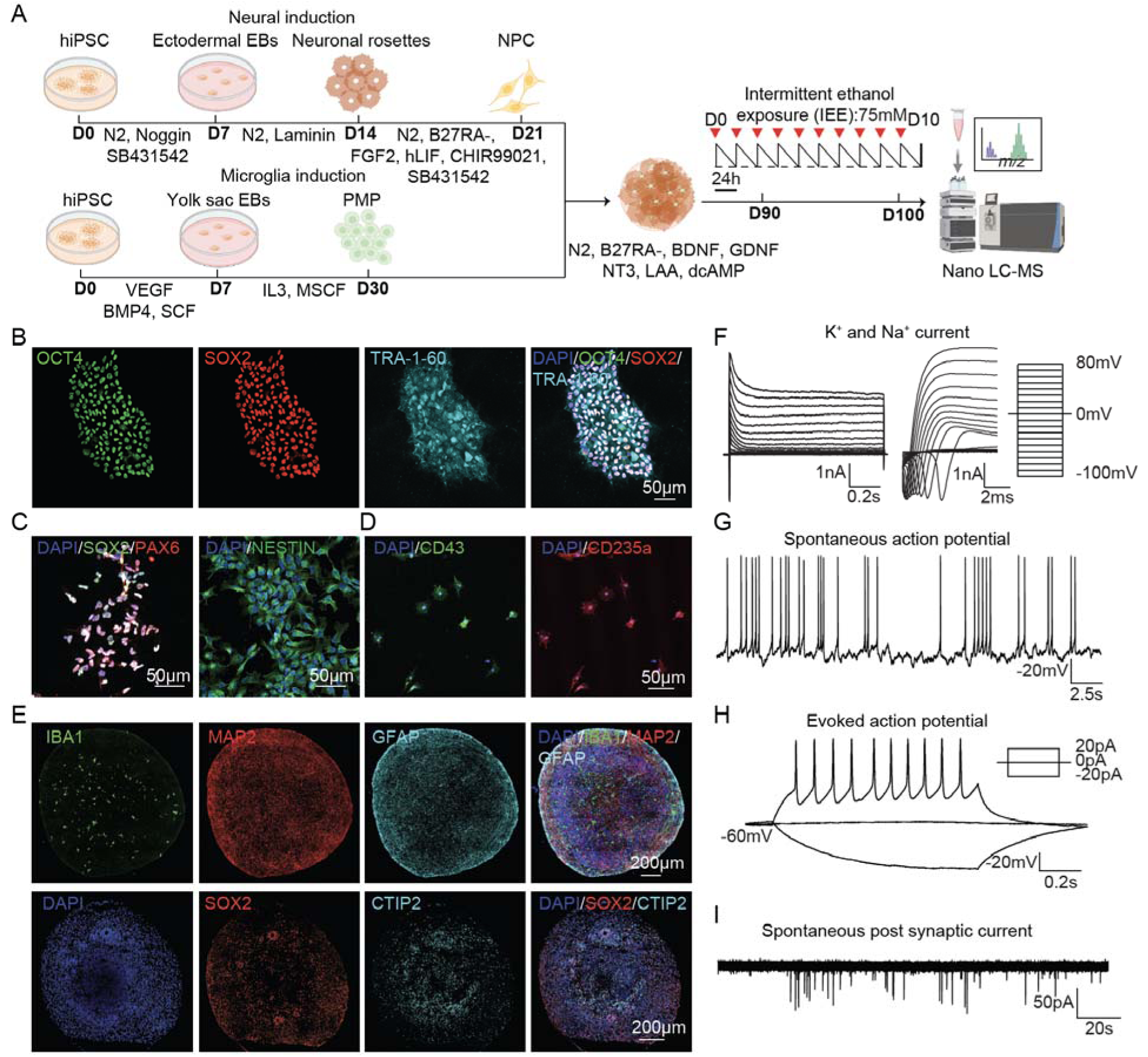
Characterization of mCOs. **(A)** Schematic of mCO generation and ethanol exposure. hiPSCs were differentiated into NPCs and PMPs, which were combined to generate mCOs. From day 90 to day 99, mCOs underwent intermittent ethanol exposure followed by LC–MS data generation. **(B)** Immunofluorescence staining of hiPSCs for the pluripotency markers OCT4, SOX2, and TRA-1-60. **(C)** Characterization of NPCs by SOX2, PAX6, and NESTIN expression. **(D)** Characterization of PMPs by CD43 and CD235a expression. **(E)** Immunofluorescence characterization of mCOs. Day-50 mCOs expressed the neural progenitor marker SOX2 and the cortical neuronal marker CTIP2. Day-100 mCOs contained IBA1-positive microglia, MAP2-positive neurons, and GFAP-positive astrocytes. **(F)** Representative whole-cell voltage-clamp recordings showing voltage-gated Na⁺ and K⁺ currents in neurons within mCOs. **(G)** Representative spontaneous action-potential firing. **(H)** Representative evoked action potentials in response to current injection. **(I)** Representative spontaneous postsynaptic currents recorded from neurons within mCOs.

### 1. Characterization of mCOs

All hiPSC lines expressed the pluripotency markers OCT4, SOX2, and TRA-1-60 (**Figure 1B**), and they have previously been shown to be euploid by eKaryotyping^30^. hiPSCs were differentiated into neuro-progenitor cells (NPCs), which expressed PAX6, SOX2, and NESTIN (**Figure 1C**). In parallel, hiPSCs were also differentiated into primitive macrophage precursors (PMPs) via yolk-sac embryonic bodies. PMPs were identified by the expression of the hematopoietic markers CD43 and CD235a (**Figure 1D**).

mCOs were generated by combining NPCs and PMPs from the same donor cell lines. These mCOs contained IBA1-positive microglia, MAP2-positive neurons, and GFAP-positive astrocytes, demonstrating the successful generation of multiple major neural cell types within mCOs (**Figure 1E**). Cortical identities of mCO neurons were demonstrated by the expression of CTIP2 (**Figure 1E**). We also performed whole-cell patch clamping on neurons in mCOs, and found they exhibited robust voltage-gated Na⁺ and K⁺ currents (**Figure 1F**), spontaneous action potential firing, and repetitive evoked action potentials in response to stepwise current injections (**Figures 1G and 1H**). These neurons also displayed spontaneous postsynaptic currents, indicating the formation of functional synaptic connections within mCOs (**Figure 1I**).

### 2. Proteomic characterization of mCOs

mCOs were intermittently exposed to either 75 mM or 0 mM ethanol for 10 days starting from day 90 followed by LC-MS. Across all mCOs, we identified a total of 131,461 unique peptides. Most peptides ranged from 7 to 20 amino acids in length, consistent with the expected peptide-length distribution following enzymatic digestion for MS–based proteomic analysis (**Figure 2A**). In total, 9,887 proteins were identified across all samples (**Figures 2B and 2C**). The distribution of missing rates is in **Figure** 1**D**, and majority of proteins had missing values in ≤6 samples (n=8,952). We also identified established markers of neurons, astrocytes, microglia, and oligodendrocyte progenitor cells, consistent with the major cell types present in mCOs (**Figure 2E**).

**Figure 2.**
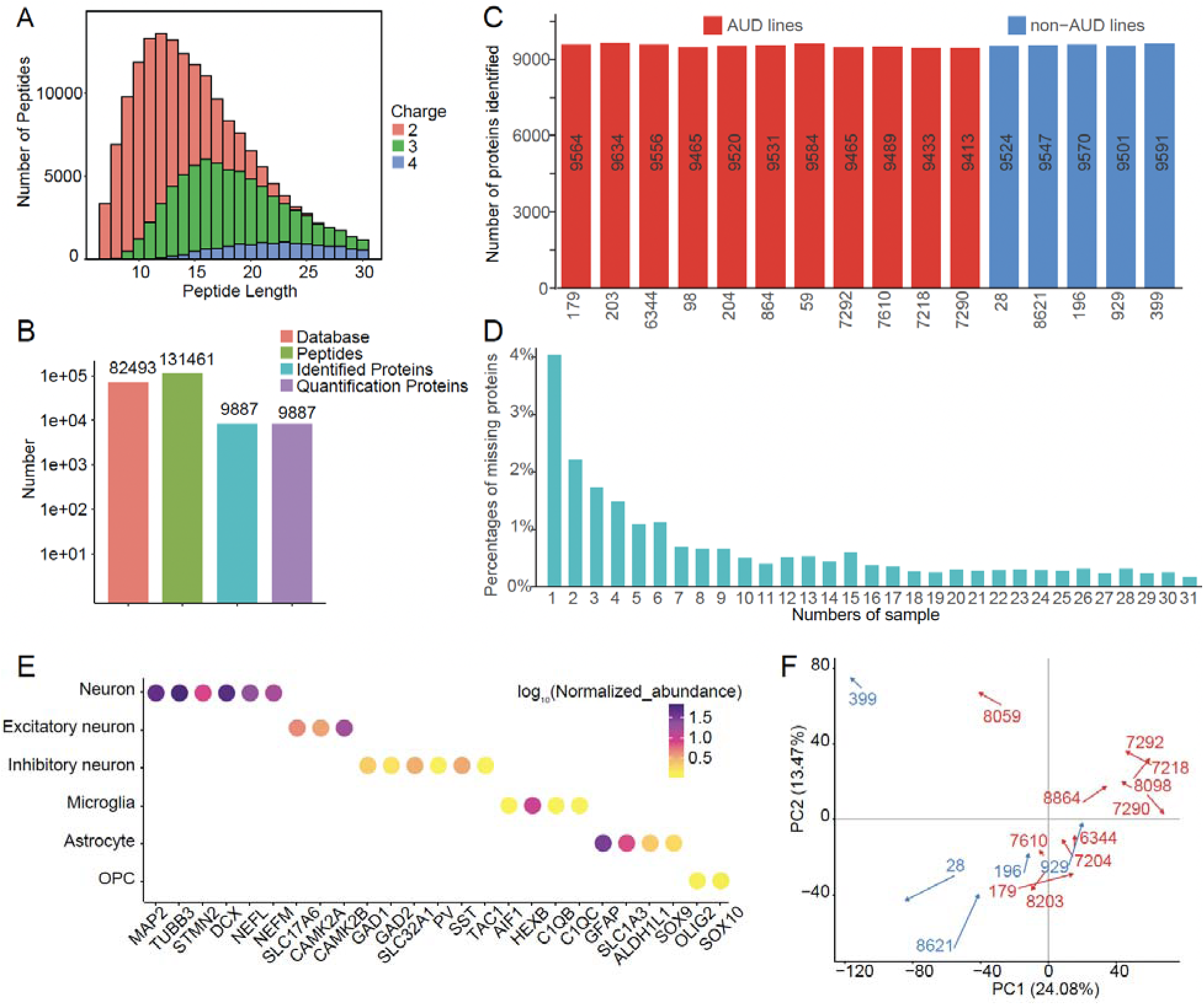
Proteomic profiling of mCOs. **(A)** Distribution of identified peptides by peptide length and charge state. **(B)** Numbers of database entries, identified peptides, identified proteins, and quantified proteins. **(C)** Number of proteins identified in each AUD and non-AUD cell line. **(D)** Distribution of protein missingness across the 32 samples. **(E)** Expression of representative protein markers for neurons, excitatory neurons, inhibitory neurons, microglia, astrocytes, and oligodendrocyte precursor cells (OPCs). Dot color indicates the average log₁₀(normalized protein abundance+1) across all cell lines. **(F)** Principal component analysis of the proteomic profiles. Arrows connect untreated and ethanol-exposed samples from the same cell line, with AUD (red) and non-AUD lines (blue) shown separately. Arrowheads indicate the direction from untreated to ethanol-exposed samples. Arrow labels are cell line IDs.

To address whether the protein expression patterns of mCOs were consistent with donors AUD status and whether ethanol exposure affects protein expressions, we performed PCA analysis and found that PC1 (24.08% variance explained) significantly separated AUD from those without AUD (P=7.8E-05), whereas PC2 (13.47% variance explained) significantly separated ethanol-treated from untreated (P=0.04) (**Figure 2F**). The separation of AUD status and ethanol exposure supported our experimental strategy for independently identifying proteins associated with inherited susceptibility and proteins responsive to ethanol exposure.

### 3. Identification of AUD-associated proteins, pathways, and candidate biomarkers

For each protein passed QC, we fitted a GLMM to test its association with AUD status while adjusting for ethanol exposure and accounting for paired samples from the same donor. GLMM models were converged for 7,591 proteins with 1,038 proteins having FDR <0.20, including 535 positively and 503 negatively associated proteins (**Supplementary Table 1**).

Reactome GSEA^40, 41^ identified 28 significantly pathways (**Supplementary Table 2**). Fifteen pathways showed positive enrichment (NES >0), indicating that they were primarily driven by proteins with higher abundance in AUD-derived mCOs. These pathways were related to neuronal and synaptic function, immune processes, and mitochondrial energy metabolism (**Figure 3A**). The remaining 13 pathways showed negative enrichment (NES <0), indicating that they were primarily driven by proteins that were less abundant in AUD-derived mCOs. These pathways were related to extracellular matrix and collagen organization, including collagen formation, laminin interactions, and extracellular matrix degradation (**Figure 3B**).

**Figure 3.**
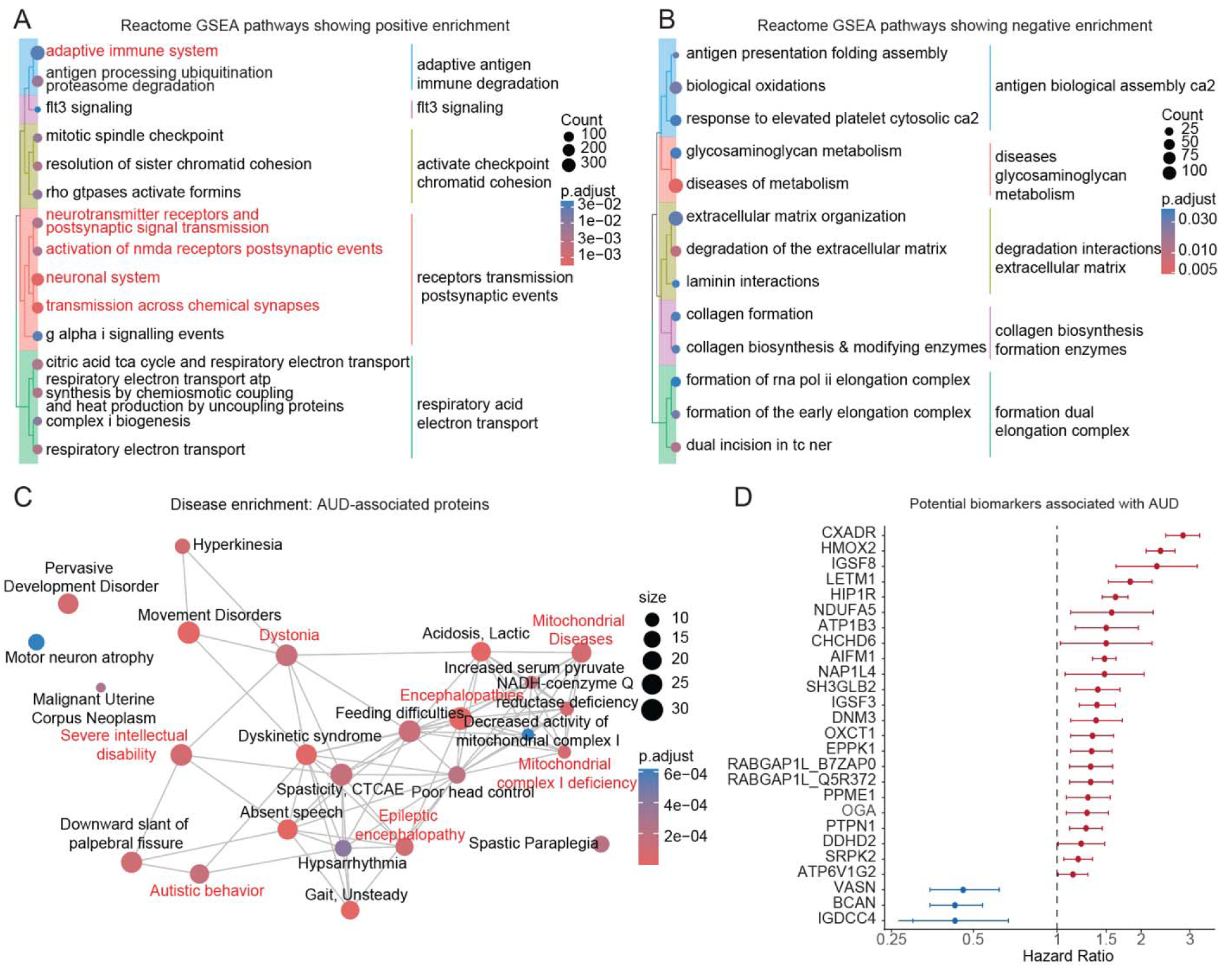
Pathway and disease enrichment analyses of AUD-associated proteins. **(A-B)** Reactome GSEA results of proteins positively (A) and negatively (B) associated with AUD, with the latter representing proteins with higher abundance in non-AUD mCOs. Dot size indicates the number of proteins, and color indicates the adjusted *P* value. Terms related to adaptive immunity, neuronal function, and synaptic transmission are highlighted in red. **(C)** Disease-enrichment network of AUD-associated proteins. Node size indicates the number of associated proteins, node color indicates the adjusted *P* value, and edges connect terms that share proteins. **(D)** Candidate AUD-associated biomarkers in UKB-PPP. Points indicate hazard ratios, and error bars indicate 95% confidence intervals. Red and blue indicate proteins positively and negatively associated with AUD, respectively.

Disease enrichment analysis further showed that proteins with abundance positively associated with AUD were enriched in 161 diseases (**Figure 3C and Supplementary Table 3)**, primarily involving neurological disorders and mitochondrial dysfunction. Representative diseases included encephalopathies, epileptic encephalopathy, dystonia, spastic paraplegia, autistic behavior, and mitochondrial complex I deficiency. In contrast, proteins negatively associated with AUD were mainly enriched in neurological, neuromuscular, and skeletal/developmental disorders **(Supplementary Table 4)**.

Among 237 proteins in both UBK-PPP incident AUD analysis and mCOs, 26 showed nominal significance (P < 0.05) and concordant effects, including 23 positively and three negatively associated proteins (**Figures 3D**; **Supplementary Table 5**). The number of concordant proteins was significantly greater than expected by chance (P =4.39 x 10⁻¹ ).

### 4. Identification of ethanol-responsive proteins, pathways, and candidate biomarkers

Paired t-tests identified 718 ethanol-responsive proteins, including 202 with increased abundance and 516 with decreased abundance following ethanol exposure (**Supplementary Table 6**).

GSEA identified 119 significant Reactome pathways (**Supplementary Table 7**). Of these, 79 showed positive enrichment, indicating increased protein abundance after ethanol exposure in these pathways. They were primarily related to DNA damage response and repair, epigenetic regulation, RNA metabolism, and protein synthesis (**Figure 4A**). Forty pathways showed negative enrichment, indicating decreased protein abundance after ethanol exposure in these pathways. They were mainly related to Intracellular vesicle trafficking, membrane transport, adaptive immune signaling, and growth factor signaling pathways (**Figure 4B**).

**Figure 4.**
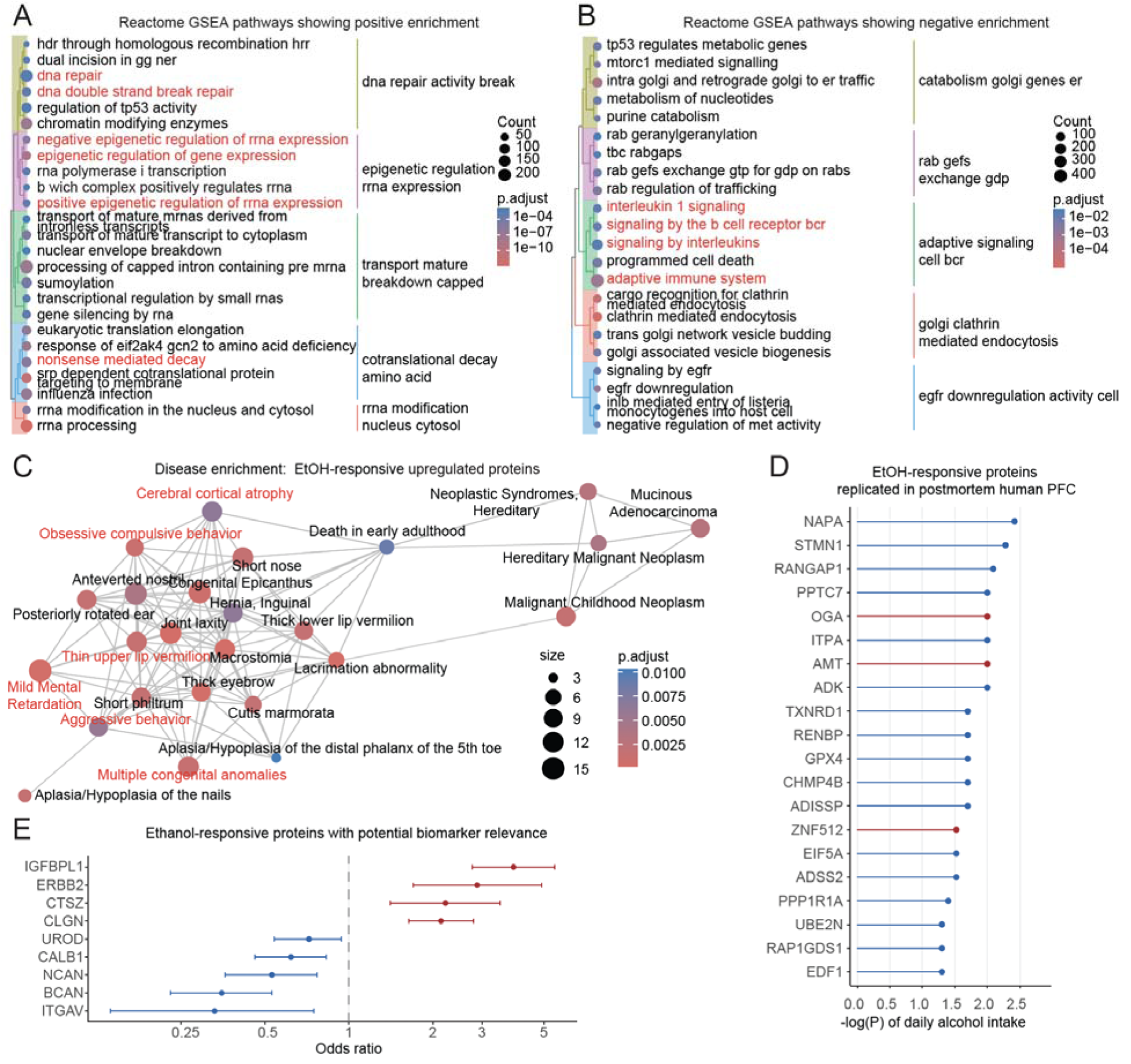
Pathway and disease enrichment analyses of ethanol-responsive proteins. **(A-B)** Reactome GSEA of proteins increased (A) or decreased (B) following ethanol exposure. Dot size indicates the number of proteins, and color indicates the adjusted *P* value. Selected terms related to DNA repair, epigenetic regulation, RNA processing, and immune signaling are highlighted in red. **(C)** Disease-enrichment network of proteins increased following ethanol exposure. Node size indicates the number of associated proteins, node color indicates the adjusted *P* value, and edges connect terms that share proteins. Red labels highlight representative pathways or disease terms. **(D)** Ethanol-responsive proteins replicated in postmortem human PFC. X-Axis is the -log(P) of daily alcohol intake; red and blue indicate increased and decreased protein abundances after ethanol exposure, respectively. **(E)** Candidate ethanol-responsive biomarker in UKB-PPP. Points indicate odds ratios, and error bars indicate 95% confidence intervals.

Disease enrichment analysis showed that proteins with increased abundance after ethanol exposure were enriched in 115 diseases primarily related to neurodevelopmental and craniofacial disorders (**Figure 4C and Supplementary Table 8**). In contrast, proteins with decreased abundance after ethanol exposure were enriched in seven disease terms related to mitochondrial and metabolic disorders, neurological diseases, and immune dysfunction (**Supplementary Table 9**).

Among 718 ethanol-responsive proteins in mCOs, 480 were detected in human postmortem PFC.^8^, of which 20 showed nominal significance (P <0.05) with concordant effects, including three with increased abundance and 17 with decreased abundance (**Figure 4D**; **Supplementary Table 10**). This overlap was greater than expected by chance (P =0.02).

Among 163 proteins in both UKB-PPP prevalent AUD analyses and mCOs, 9 showed nominal significance (P <0.05) and concordant effects, including four with increased abundance and five with decreased abundance (**Figure 4E**; **Supplementary Table 11**). The number of concordant proteins was significantly greater than expected by chance (P =0.02).

### 5. Differential ethanol responses in AUD and non-AUD mCOs using WGCNA

We observed limited overlap between AUD-associated proteins and ethanol-responsive proteins, with only 9 proteins showing concordant increases and 39 showing concordant decreases (**Supplementary Table 12**). This indicates that baseline AUD-associated differences and ethanol-induced changes involve largely distinct protein sets. We therefore investigated whether the proteomic response to ethanol differed between AUD and non-AUD mCOs using WGCNA and linear mixed-effects models. Modules identified by WGCNA are shown in **Figure 5A**, of which 17 were significantly altered by ethanol exposure (**Figure 5B).** After fitting linear mixed-effects models, we found different ethanol responses between AUD and non-AUD mCOs. The black module increased after ethanol exposure mainly in AUD mCOs. Reactome enrichment analysis showed that this module was associated with RNA metabolism and processing, including mRNA processing, RNA catabolism, RNA transport, DNA damage response, and chromatin and chromosome organization pathways (**Figure 5C**). The yellow module decreased in both AUD and non-AUD mCOs, but the decrease was greater in non-AUD mCOs after ethanol exposure (**Figure 5B**). This module was enriched in intracellular protein trafficking, membrane transport, neuronal signaling, and lipid metabolism pathways, suggesting that ethanol disrupts protein transport and membrane homeostasis (**Figure 5D**).

**Figure 5.**
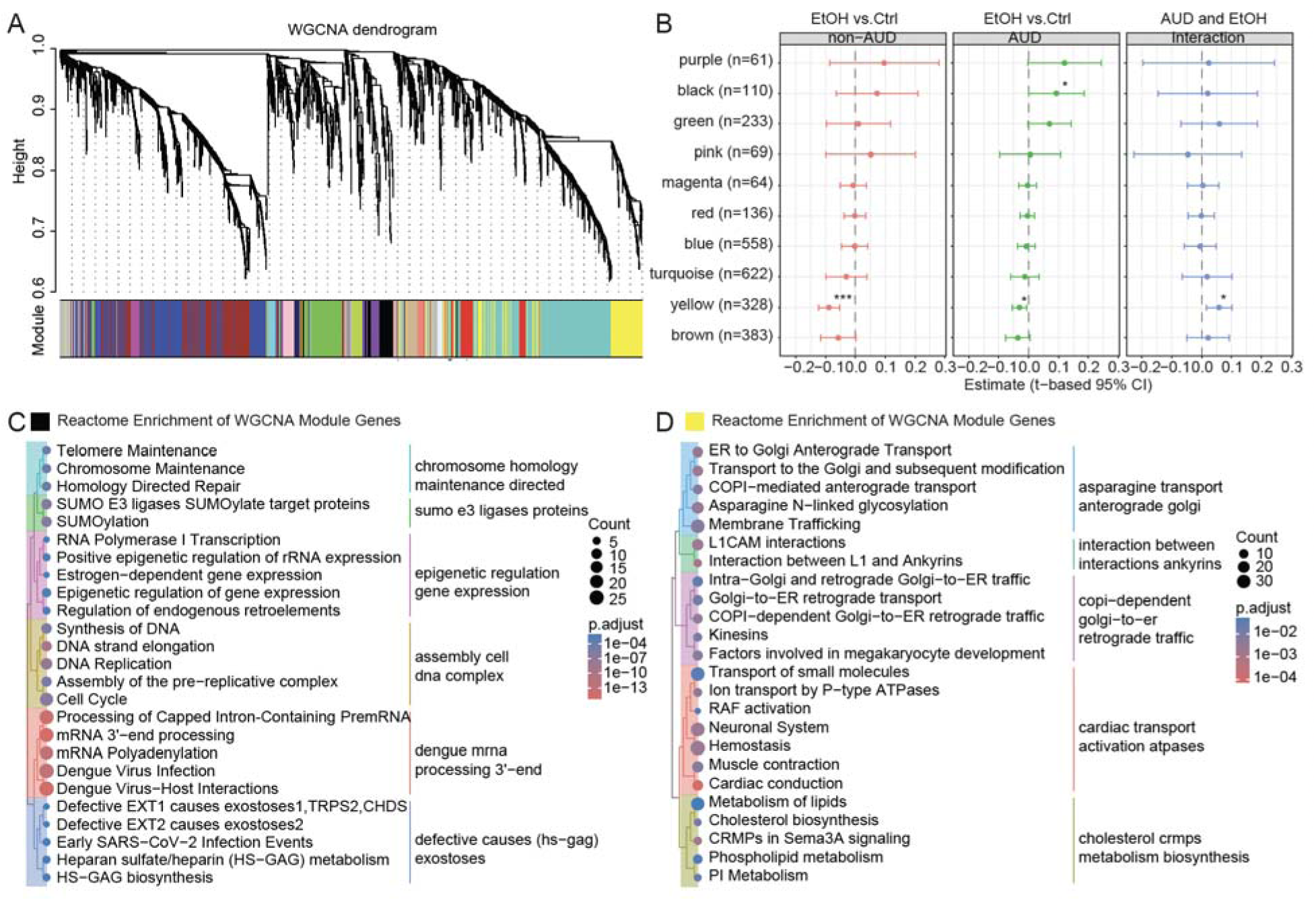
WGCNA identifies protein modules with differential ethanol responses in AUD and non-AUD mCOs. **(A)** WGCNA dendrogram showing protein co-expression modules, with module assignments indicated by the color bar. **(B)** Estimated ethanol effects on module eigengenes in non-AUD and AUD mCOss and the AUD-by-ethanol interaction. Points indicate model estimates, and error bars indicate 95% confidence intervals. *p<0.05, ***p<0.001. **(C)** Reactome enrichment analysis of proteins in the Black module, which increased after ethanol exposure mainly in AUD mCOs. Dot size indicates the number of proteins, and color indicates the adjusted *P* value. **(D)** Reactome enrichment analysis of proteins in the Yellow module, which decreased more strongly after ethanol exposure in non-AUD mCOs. Dot size indicates the number of proteins, and color indicates the adjusted *P* value.

### 6. Polygenic scores (PGS) are associated with major AUD-related proteomic variation

Since donors were selected also based on AUD PGS, we tested correlations between the first two principal components (PC1 and PC2; **Figure 2F**) of mCO proteome and PGS. Pearson correlation analyses showed that PC1, which primarily distinguished cell lines by AUD status, was also significantly correlated with both original and latest PGS (**Figures 6A-B, E)**, whereas PC2, which was linked to ethanol exposure, showed no association with either PGS (**Figure 6C**)

**Figure 6.**
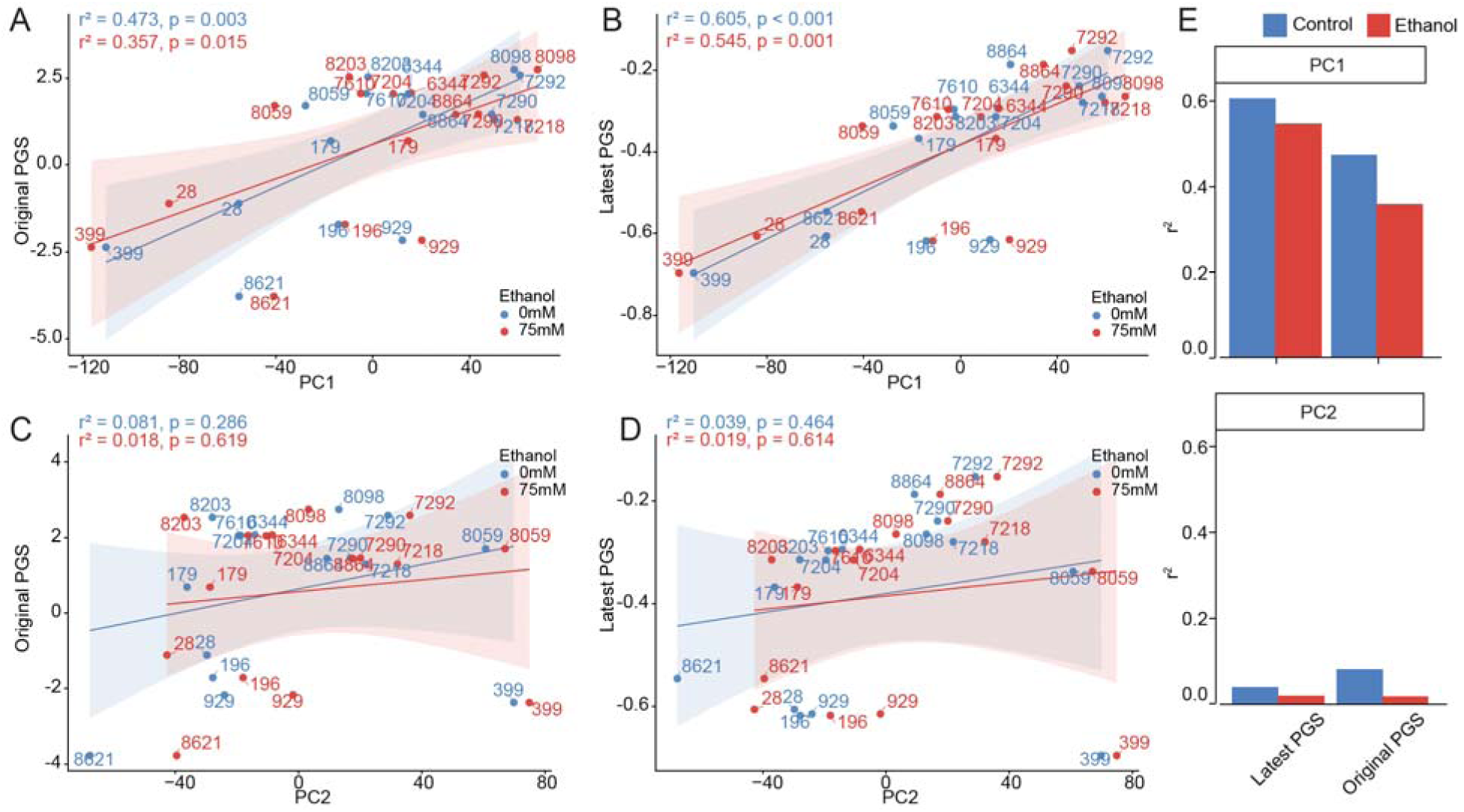
Associations between polygenic scores and major proteomic variation. (**A-B**) Pearson correlations between PC1 and polygenic scores (PGS): Latest PGS (**A**), Original PGS (**B**), shown separately for control (0 mM, blue) and ethanol-exposed (75 mM, red) samples. (**C-D**) Pearson correlations between PC2 and polygenic scores (PGS): Latest PGS (**C**), Original PGS (**D**), shown separately for control (0 mM, blue) and ethanol-exposed (75 mM, red) samples. Lines indicate linear fits with shaded 95% confidence intervals. Pearson correlation (r^2^) and P values are shown for each condition. Numbers indicate individual cell lines. (**E**) Comparison of r² values between each PGS and PC1 or PC2 in control (blue) and ethanol-exposed (red) samples.

## Discussion

In this study, we used hiPSC-derived, microglia-containing cortical organoids generated from donors with/without AUD and high/low PGS together with a donor-matched intermittent ethanol exposure design to identify AUD-associated and ethanol-responsive proteins. Enrichment analyses demonstrated that these two types of proteins were enriched in different pathways.

Twenty ethanol-responsive proteins were replicated in an independent proteomic dataset from human postmortem PFC. Integration with UKB-PPP plasma proteomic data identified 26 candidate AUD-associated biomarkers and 9 candidate ethanol-responsive biomarkers. Finally, PC1 of mCO proteome explained ∼60% of PGS calculated using the largest GWAS of PAU.

As demonstrated in our study, AUD-associated and ethanol-responsive proteins showed limited overlap (**Supplementary Table 12**), suggesting that inherited susceptibility to AUD and ethanol exposure may predominantly influence distinct biological processes. Ideally, these two types of proteins would be identified directly from human brain tissues. However, brain tissues cannot be prospectively obtained from individuals who may later develop AUD, making it particularly challenging to identify AUD-associated proteins. Our iPSC-derived, microglia-containing organoid model provides an opportunity to address this challenge. By including donors with differing AUD status and comparing untreated and ethanol-exposed mCOs from the same donor, we were able to examine baseline molecular differences associated with AUD susceptibility and direct proteomic responses to ethanol within the same experimental system. Importantly, the selection of donors with extreme PGS further enabled us to enrich for differences in inherited liability to AUD. Moreover, this strategy may potentially be extended to human postmortem brain studies, where comparison of individuals with extremely high versus low AUD PGS could help identify molecular signatures associated with genetic liability for AUD. Together, our design provides a framework for disentangling molecular signatures associated with predisposition to AUD from those induced by ethanol exposure.

AUD-associated proteins were mainly enriched in two biological processes. First, proteins with higher abundance in AUD-derived mCOs were enriched in neuronal signaling and synaptic transmission. Selected examples of pathways affected by this group include core components of the SNARE machinery (e.g., VAMP2, SNAP25), synaptic plasticity mechanisms (e.g., CAMK2A, PRKACA), and membrane excitability components (e.g., CACNA1E, KCNA6).

Disease enrichment similarly identified neurological and neurobehavioral terms, including encephalopathies, dystonia, and autistic behavior. These findings are consistent with evidence that AUD is related to the remodeling of glutamatergic signaling, including alterations in glutamate receptor function and synaptic plasticity^53^. Altered glutamatergic signaling plays a central role in the development and maintenance of alcohol-related behaviors such as alcohol seeking and dependence^54, 55^. Second, AUD-associated proteins were also enriched in respiratory electron transport and oxidative phosphorylation pathways, and disease enrichment highlighted multiple mitochondrial disorders. Among proteins linked with these functions are several subunits of electron transport Complexes I, III, and V as well as members of the pyruvate dehydrogenase complex and the TCA cycle. These findings further support the involvement of mitochondrial pathways in AUD shown in previous studies^56, 57^ ^58^. Together, our findings suggest that altered neuronal signaling and mitochondrial energy metabolism are key molecular features associated with AUD.

In contrast, ethanol exposure mainly changed abundance of proteins involved in epigenetic regulation, DNA repair, and RNA processing. These pathways were also enriched in the WGCNA module that exhibited a greater difference after ethanol exposure in mCOs from those with AUD compared to those without. Previous studies have shown that alcohol consumption was associated with widespread DNA damage and methylation changes^59–61^, e.g., in mouse primary cortical neurons, prolonged ethanol metabolism damaged DNA and caused inaccurate DNA repair, abnormal cell-cycle activation, and a senescence-like state^61^. In developing human cortical tissue, ethanol exposure induced hundreds of alternative splicing events in genes involved in cell death, cell junctions, and synapse formation^62, 63^. Interestingly, genetically regulated alternative splicing has also been implicated in AUD susceptibility, with alcohol-related splicing events linked to neurogenesis and gliogenesis pathways^64^. Consistent with these molecular alterations, our disease enrichment analysis highlighted neurodevelopmental, neurobehavioral, and craniofacial disorders, potentially reflecting shared proteins involved in nervous system development, RNA processing, DNA repair, and genome maintenance.

Several identified candidate biomarkers were supported across multiple datasets. BCAN was negatively associated with AUD, decreased expression after ethanol exposure, and showed concordant associations in UKB-PPP incident and prevalent AUD analyses and an Icelandic plasma cohort^24^. Although BCAN is mainly involved in extracellular matrix organization and has been linked to neuroimaging traits^65^, proteomic studies have also implicated it in depression, autism spectrum disorder, and posttraumatic stress disorder^66–68^, suggesting broader relevance across psychiatric conditions. OGA, a regulator of protein O-GlcNAcylation, was both AUD-associated and ethanol-responsive, replicated in postmortem PFC data, and was associated with incident AUD in UKB-PPP. It has also been linked to several psychiatric and neurological traits, such as attention deficit-hyperactivity disorder, autism spectrum disorder, cognitive function, epilepsy, and neuroticism^69^. RABGAP1L showed positive associations with AUD for both the full-length protein (Q5R372) and an isoform (B7ZAP0) and was associated with incident AUD in UKB-PPP, and interestingly, it had previously been implicated by a COGA GWAS^70^ and subsequent GWAS of alcohol-related phenotypes^6, 71^. STMN1 (stathmin 1) expression increased after ethanol exposure, was replicated in postmortem PFC data and associated with prevalent AUD in UKB-PPP, and has been linked to alcohol consumption^71^.

Additionally, CXADR (CXADR cell adhesion molecule), VASN (vasorin), and IGDCC4 (immunoglobulin superfamily DCC subclass member 4) were AUD-associated and were replicated in UKB-PPP incident AUD and an Icelandic plasma study^24^. These AUD-associated and/or ethanol-responsive proteins, detectable in plasma, represent candidate biomarkers that require further investigation.

Correlations of PC1 with PGS indicated that PC1 explained approximately 40–60% of the variance in PGS. Given the intentional selection of donors based on AUD status and extreme PGS, this finding raises the possibility that PC1 captures substantial AUD-related variation, although the magnitude of the associations should be interpreted in the context of this enriched sample. Nevertheless, the proportion of PGS variance explained by PC1 is notably similar to the magnitude of estimated AUD heritability (50–60%)^51, 72, 73^. Although these quantities are not directly comparable, their similar magnitude raises the possibility that the proteomic variation captured by PC1 reflects a substantial component of the genetic liability to AUD. Because the predictive performance of PGS varies according to the discovery GWAS^74, 75^, we recalculated PGS using the largest PAU GWAS^51^. Associations with PC1 were largely consistent between these two PGS, with the PGS derived from the largest PAU GWAS showing stronger correlation (**Figure 6B**). Our findings, while preliminary, suggest that continued improvements in PGS may enhance their utility for selecting informative samples, thereby increasing statistical power and reducing the cost of studies aimed at elucidating molecular mechanisms underlying AUD.

While our study is the first to identify distinct AUD-associated and ethanol-responsive proteins and biomarkers in a human microglia-containing cortical organoid model, there are some limitations. First, although selecting donors with AUD/high PGS and controls/low PGS and using untreated and ethanol-exposed mCOs from each donor improved statistical power, the sample size remained relatively modest because of the labor-intensive and costly nature of human stem cell preparation, brain organoid generation, and deep proteomic profiling. Second, mCOs, while providing a simplified model for investigation, do not fully recapitulate the cellular diversity, vascularization, immune environment, or long-term maturation of the adult human brain. Third, all donors were of European ancestry, and replication in more diverse populations will be important to evaluate the generalizability of these findings. Fourth, replication in postmortem human brain tissue was constrained by the limited sample size and substantially lower proteome coverage of the available dataset, reducing the number of proteins that could be evaluated. Finally, the present study identified proteins associated with inherited AUD susceptibility and ethanol exposure but cannot establish their causal roles in disease pathogenesis. Future studies combining CRISPR-mediated gene editing and pharmacological perturbation will help determine the functional roles of these candidate proteins and pathways.

In conclusion, this study provides a comprehensive proteomic profile of microglia-containing cortical organoids from donors with/without AUD and high/low PGS under untreated and ethanol-exposed conditions. Our findings help distinguish proteomic alterations associated with inherited susceptibility to AUD from those induced by ethanol exposure, identify candidate biomarkers, and demonstrate a strong relationship between major proteomic variation and polygenic liability for AUD. More broadly, our study supports the utility of microglia-containing cortical organoids stratified by AUD status and PGS as a novel platform for investigating the molecular mechanisms underlying AUD and provides a useful resource for prioritizing proteins and pathways for future functional and biomarker studies.

## Acknowledgements

The Collaborative Study on the Genetics of Alcoholism (COGA), Principal Investigators B. Porjesz, V. Hesselbrock, A. Agrawal; Scientific Director, A. Agrawal; Translational Director, D. Dick, includes nine different centers: University of Connecticut (V. Hesselbrock); Indiana University (H.J. Edenberg, T. Foroud, Y. Liu, M.H. Plawecki); University of Iowa Carver College of Medicine (A. Andersen S. Kuperman); SUNY Downstate Health Sciences University (B. Porjesz, J. Meyers); Washington University in St. Louis (L. Bierut, A. Agrawal, S. Hartz); University of California at San Diego (M. Schuckit); Rutgers University (D. Dick, R. Hart, J. Salvatore, J. Tischfield); The Children’s Hospital of Philadelphia, University of Pennsylvania (L. Almasy); Icahn School of Medicine at Mount Sinai (A. Goate, P. Slesinger); and Howard University (D. Scott). Other COGA collaborators include: M. Hesselbrock, K. Manning (University of Connecticut); D. Lai, J. Nurnberger Jr., L. Wetherill, A. Miller, X. Xuei, (Indiana University); J. Kramer (University of Iowa), G. Chan (University of Iowa; University of Connecticut); C. Kamarajan, A. Pandey, D.B. Chorlian, P. Barr, S. Kinreich, G. Pandey, Z. Neale, C. Chatzinakos, J. Zhang, S. Saenz deViteri, A. Bingly (SUNY Downstate); G. Pathak (Icahn School of Medicine at Mount Sinai); A. Anokhin, K. Bucholz, F. Dong, A. Hatoum, E. Johnson, J. Rice, S. Saccone (Washington University); F. Aliev, Z. Pang, S. Kuo, S. Brislin, (Rutgers University). We continue to be inspired by our memories of Henri Begleiter and Theodore Reich, founding PI and Co-PI of COGA, and also owe a debt of gratitude to other past organizers of COGA, including Ting-Kai Li, P. Michael Conneally, Raymond Crowe, and Wendy Reich, for their critical contributions. This national collaborative study is supported by NIH Grant U10AA008401 from the National Institute on Alcohol Abuse and Alcoholism (NIAAA) and the National Institute on Drug Abuse (NIDA).

Y. Liu and D. Lai are supported by National Institutes of Health award number AA031176.

X. Li, R. Hart, and P. Pang are supported by National Institutes of Health award number R01AA023797.

A.J. Boreland is supported by NINDS T32NS115700.

